# High-Frequency Spinal Cord Stimulation Reorganizes Cortical Cross-Frequency Coupling in a Region- and Time-Dependent Manner

**DOI:** 10.64898/2025.12.15.694369

**Authors:** Vishal Bharmauria, Hiroyuki Oya, Yarema Bezchlibnyk, Nour Shaheen, Amirhossein Ghaderi, Yahia Yassine Belkacemi, Karim Johari, Arun Singh, Alexander L Green, Hiroto Kawasaki, Can Sarica, Brian Dalm, Andres M Lozano, Matthew A Howard, Oliver Flouty

## Abstract

Pain management strategies have progressed beyond traditional pharmacologic and physical interventions, integrating advanced neuromodulation techniques such as deep brain stimulation, peripheral nerve stimulation, and high-frequency spinal cord stimulation (hSCS). Despite its clinical efficacy, the supraspinal mechanisms underlying hSCS remain poorly understood. Prior work in sheep demonstrated that hSCS modulates gamma (γ) band (70–150 Hz) activity in the primary somatosensory and association cortices, implicating cortical involvement in pain modulation. Given, the interaction between low and high oscillations, we hypothesized that hSCS modulates γ activity in a region- and time-dependent manner through specific coupling with theta (□) rhythms (4–8 Hz). Using 96-channel subdural electrocorticography (ECoG), we computed □-γ phase–amplitude coupling (PAC) and the corresponding modulation index (MI) to quantify the effects of hSCS. While the preferred □phase of γ activity remained consistent across conditions and regions, MI increased significantly post-stimulation—most prominently in the association cortex, where robust -γ phase locking was observed. In contrast, the somatosensory cortex exhibited weaker and more variable locking. Temporally, both cortices demonstrated an early, rapid increase in MI post-hSCS, accompanied by a shift (association) and attenuation (somatosensory) of the secondary peak. These findings reveal distinct regional and temporal dynamics in PAC following hSCS and suggest complementary roles of somatosensory and association cortices in processing neuromodulatory input. hSCS appears to reorganize cortical cross-frequency interactions, supporting its role in reorganizing functional network dynamics relevant to sensory processing and the subjective pain experience.

## INTRODUCTION

Pain management encompasses several approaches, including medications, physical therapies, natural remedies, and more recently, simulation-based neuromodulation techniques (Johnson et al., 2013; Chakravarthy et al., 2016; Flouty et al., 2022; Schulder et al., 2023; Shaheen et al., 2023; Shaheen and Flouty, 2024; Sun et al., 2024; Wang and Doan, 2024; Mogedano-Cruz et al., 2025; Puppalla et al., 2025). Traditionally, paresthesia-based spinal cord stimulation (P-SCS), where electrical impulses generating tingling or buzzing sensation to mask pain, was commonly used for pain relief (Johnson et al., 2013; Hagedorn et al., 2021). Nowadays, stimulation-based techniques have evolved into neuromodulation strategies such as dorsal root ganglion stimulation (DRG-S), burst SCS (B-SCS), and high-frequency SCS (hSCS), each targeting distinct neuroanatomical substrates to enhance pain relief (De Ridder et al., 2013; Kapural et al., 2015; Caylor et al., 2019; Kirketeig et al., 2019; Abraham et al., 2021; Ehsanian et al., 2024). While these approaches offer improved clinical outcomes, a unified mechanistic understanding—particularly regarding their supraspinal effects—remains elusive. Previous work in sheep demonstrated : 1) increased high-frequency (γ) amplitude as a function of dorsal column stimulation voltage (Flouty et al., 2013) and 2) hSCS robustly suppressed high-frequency (γ) oscillations in both primary somatosensory and association cortices, implicating oscillations involved in local cortical processing as a key modulatory target (Bharmauria et al., 2025).

Oscillatory activity is essential for coordinating brain function across spatial and temporal scales (Buzsáki, 2006; Fries, 2015). High-frequency γ oscillations (70–150 Hz) are associated with local circuit computations, while low-frequency rhythms such as (4–8 Hz) have been shown to mediate long-range coordination and top-down control (Gross et al., 2007; Wang et al., 2011; Lisman and Jensen, 2013; Taesler and Rose, 2016; Song et al., 2025). Importantly, recent studies suggest that pain alters the balance and interaction between these frequencies, particularly via phase–amplitude coupling (PAC), a mechanism whereby the amplitude of high-frequency activity is modulated by the phase of slower rhythms (Oshiro et al., 2009; Wang et al., 2011, 2016; Ploner et al., 2017; Ong et al., 2019; Tan and Kuner, 2021; Kong et al., 2024; Song et al., 2025). Aberrant PAC has been reported in several pain-related cortical regions, including the insula, anterior cingulate, and prefrontal cortex, and may reflect disrupted network integration in chronic pain (Oshiro et al., 2009; Wang et al., 2016; Ploner et al., 2017; Ong et al., 2019; Tan and Kuner, 2021).

Building on previous findings (Flouty et al., 2012, 2013; Huang et al., 2014; Bharmauria et al., 2025), this study tests the hypothesis that hSCS modulates cortical sensory processing by disrupting abnormal □-γ coupling across somatosensory and association cortices, regions that are reciprocally connected with each other (Pandya and Yeterian, 1985; Kaas, 1993, 2004). By employing subdural electrocorticography (ECoG) in sheep to quantify PAC before and after hSCS, we assessed how the strength of □-γ coupling, quantified as modulation index (MI), was altered by hSCS. We demonstrate that: 1) overall, after hSCS - □ γ phase-amplitude locking (preferred phase angle with high □γ) remained largely unchanged across both cortices 2) MI exhibited predominantly an increase, following stimulation. 3) Temporally, a reciprocal, complementary, shift in MI between the two cortices emerged after hSCS, suggesting a dynamic redistribution of coupling strength. Briefly, this study highlights the importance of □-γ coupling in maintaining the network organization for sensory perception while regulating the amplitude gain of responses.

## METHODS

### Ethics Approval

All experimental procedures were approved by the University of Iowa Animal Care and Use Committee (IACUC#0902039). Four adult sheep (Ovis aries; 58–71 kg) were used for this study.

### Surgical Preparation and Electrode Placement

Sheep were anesthetized with Isoflurane and maintained under general anesthesia via endotracheal intubation. Prior to incision, 10–20 cc of 2% Lidocaine was administered locally. A longitudinal incision was made near the left calcaneus to expose the tibial nerve for stimulation **(Figure 1Ai)** where two EMG electrodes were inserted. A right-sided hemi-craniectomy was performed to expose the cerebral cortex. A 96-channel subdural grid (Ad-Tech Medical; 1.4 mm contact diameter, 3.0 mm inter-contact spacing) was placed over the somatosensory and association cortices **(Figure 1Aii)**. These cortical regions are reciprocally interconnected and play distinct roles in sensory integration and higher-order processing (Pandya and Yeterian, 1985).

**Figure 1.**
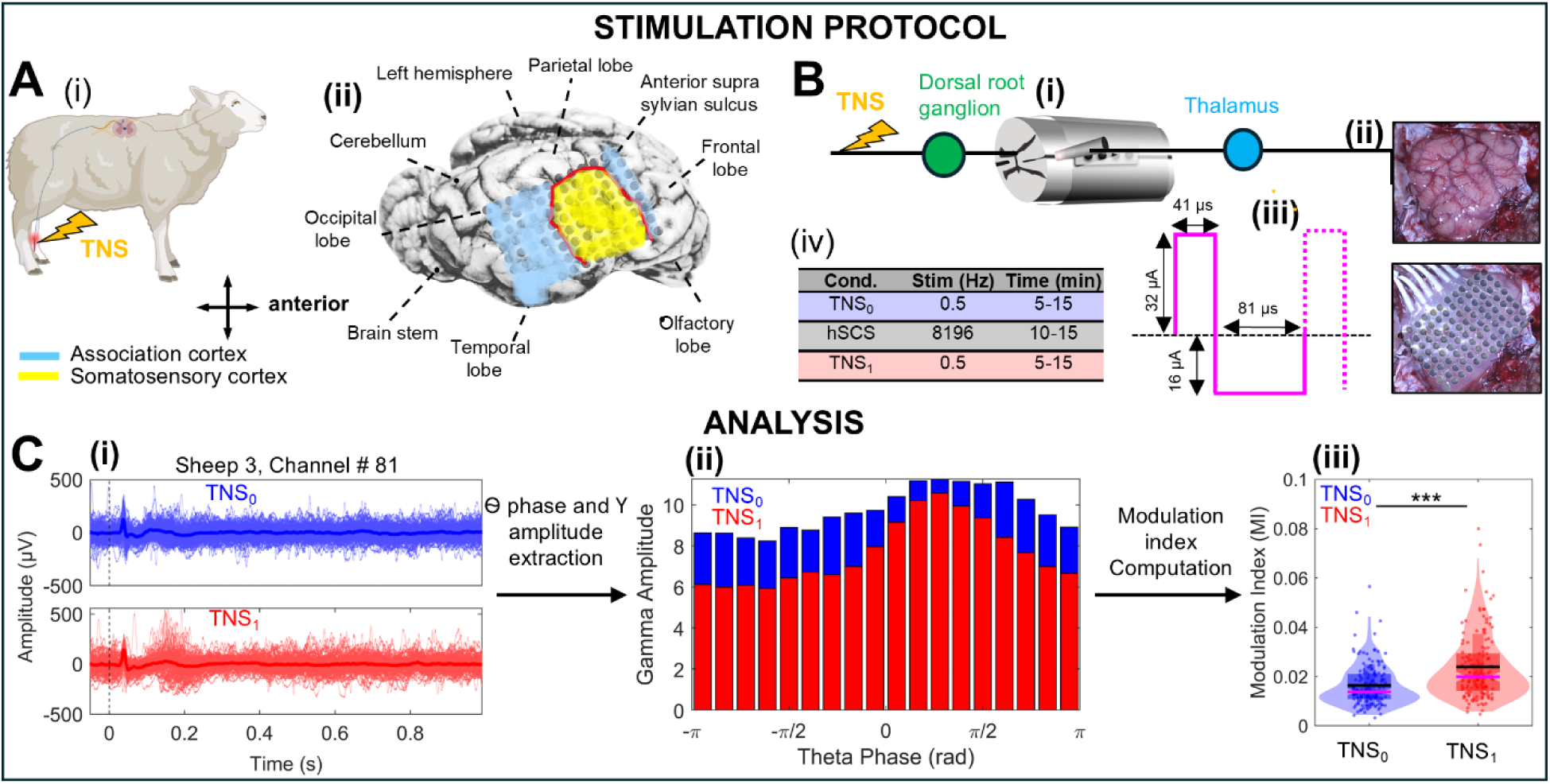
Experimental setup and analysis pipline. **(A)** ECoG placement in sheep brain during TNS **(i)** Schematic of the mammalian sensory pathway illustrating information flow from the periphery to higher order cortical areas. **(ii)** Dorsal view of a sheep brain implanted with a 96-contact subdural ECoG grid spanning the somatosensory cortex (highlighted in yellow) and association cortex (highlighted in blue). **(B)** Stimulation protocol and timeline**. (i)** Analysis pipeline High-frequency spinal cord stimulator (hSCS) used for delivering stimulation. **(ii)** Photograph of the exposed cortical surface without (top) and with (bottom) the grid in Sheep 1, providing anatomical reference. **(iii)** Summary of the stimulation parameters used during the experiment, including frequency, pulse width, and amplitude. **(iv)** Timeline outlining the experimental protocol, including pre-stimulation (TNS_0_), stimulation, and post-stimulation (TNS_1_) recording periods. **(C)** Analysis Pipeline **(i)** Evoked potential aligned to TNS onset (black broken line) for all trials (Sheep 3, channel 81) in TNS_0_ (blue) and TNS_1_ (red) condition. **(ii)** -γ locking at TNS_0_ (blue) and TNS_1_ (red) conditions**. (iii)** Modulation index computation. For channel 81 in Sheep 3, a significant increase was noticed between conditions (Mann Whitney U test, P < 0.001)

To deliver hSCS, a laminectomy was performed from thoracic levels T7–T10. The thecal sac was incised open under the surgical microscope exposing the dorsal spinal cord. A custom bipolar stimulation probe was positioned on the spinal surface over T8-T9 using a precision micromanipulator to ensure accurate placement **(Figure 1Bi)**. A separate subdural strip electrode was placed near the spinal cord to monitor local field potentials.

The exposed spinal cord was then irrigated and flooded with saline. During electrophysiological recordings from ECoG channels on the cortex (**Figure 1Bii)**, anesthesia was transitioned to a low-dose Propofol infusion (0.4 mg/kg/hr) to minimize its suppressive effects on cortical oscillations. Anesthesia depth was monitored using corneal reflex, and core body temperature was maintained with a heating pad. After completion of the experiment, animals were euthanized using intravenous pentobarbital (120 mg/kg).

### Experimental Paradigm and Stimulation Protocol

Before stimulation, a 30-minute rest period was given for anesthetic washout. The total surgical and experimental time was 13 ± 2 hours. Each animal underwent three stimulation blocks (**Fig. 1Biii-iv)**. The experimental sequence consisted of: (1) tibial nerve stimulation alone (TNS_0_), (2) combined tibial nerve and hSCS, and (3) a second block of tibial nerve stimulation alone (TNS_1_) **(Fig. 1Biv)**. This design allowed us to isolate the cortical effects of hSCS by comparing evoked neural responses before and after stimulation.

#### TNS stimulation

Tibial nerve stimulation (TNS) was administered using a Grass constant current stimulator at 0.5 Hz (one pulse every 2 seconds) with a pulse width of 200 µs. This protocol reliably evoked potentials across trials, allowing for trial-by-trial analysis of phase-amplitude coupling (PAC) metrics **(Fig. 1Biii).** To ensure that the responses were not influenced by sensitization or desensitization, we evaluated their stability by comparing mean responses across early, middle, and late trials during the TNS_0_ condition in all sheep. No evidence of response modulation over time was observed (Bharmauria et al., 2025).

#### hSCS Stimulation

High-frequency spinal cord stimulation (hSCS) was applied using a custom bipolar probe with two spherical contacts (1.0 mm diameter, 2.0 mm spacing), delivering constant-current, charge-balanced square pulses (122 µs duration) at 8.196 kHz via a Tucker-Davis Technologies IZ-2 stimulator. Stimulation duration varied across animals (5–15 minutes).

### Electrocorticography (ECoG) and Signal Acquisition

ECoG data were recorded using a 96-channel grid connected to a Tucker-Davis Technologies RZ-2/PZ-2 acquisition system at a 5 kHz sampling rate. Signals were visually inspected and notch-filtered at 60 Hz to remove line noise.

### Signal Processing and Theta–Gamma (□-**γ**) Analysis

All analyses were performed offline using custom MATLAB scripts. Noisy trials were excluded if their peak amplitude exceeded 10 standard deviations from the mean. Time-locked epochs were extracted from -0.05 to 1 sec post-TNS onset for each trial.

To quantify PAC between phase and γ amplitude, we computed the instantaneous phase of the band and the amplitude envelope of the γ band using MATLAB’s signal processing functions. For each channel and trial, the time series was bandpass-filtered separately in the (4–8 Hz) and high-γ (70–150 Hz) frequency bands using the *‘bandpass’* function, which applies zero-phase digital filtering (via *‘filtfilt’ function*) to avoid phase distortion. The analytic signal of the filtered data was then derived using the *‘hilbert’* function, which computes the Hilbert transform to return a complex-valued signal. From this analytic signal, the instantaneous phase was extracted using the *‘angle’* function, and the γ amplitude envelope was obtained as the magnitude of the analytic signal using *‘abs’*.

These computations provided time-resolved phase and amplitude values for each trial and channel. To assess how γ amplitude was modulated by phase, we aggregated the instantaneous values across all time points and trials. The phase range (−π to π radians) was divided into 18 equal-sized bins, and the mean high-γ amplitude was calculated within each bin. This yielded a phase-amplitude profile for each channel, showing how γ amplitude varied with phase [i.e., theta-high gamma (□-γ) coupling, reflected in the preferential phase where γ amplitude was most concentrated] for the *TNS_0_*and *TNS_1_* conditions. Phase locking, which quantifies the consistency of γ amplitude occurring at a specific phase, was measured using the mean resultant length (MRL), ranging from 0 (γ amplitude uniformly distributed across phases, no locking) to 1 (γ amplitude consistently aligned to a preferred phase, perfect locking). Phase locking captures the alignment of fast oscillatory activity to slower rhythms and complements the modulation index (MI; see below), which quantifies the strength of□–γ PAC. Together, these metrics provide a comprehensive assessment of how hSCS modulates cross-frequency interactions in the somatosensory and association cortices.

### Modulation index computation

To quantify the strength of -γ PAC, a modulation index (MI) was calculated following the method described by Tort and colleagues (Tort et al., 2008, 2010). For each trial, the phase–amplitude distribution was computed and normalized to yield a probability distribution of high-γ amplitude across phase bins. The Shannon entropy of this empirical distribution was then compared with the entropy of a uniform distribution using the following formula:

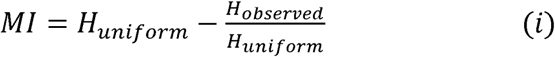

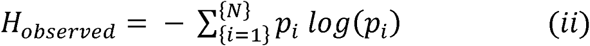

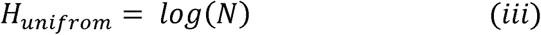

where, *H_observed_* is the Shannon entropy of the empirical amplitude distribution, *H_uniform_* is the entropy of the distribution over N bins (18 in this case), and *p_i_* is the normalized mean amplitude (probability) in the *i^th^* phase bin.

This normalized Kullback–Leibler divergence yields an MI value between 0 (no coupling, the gamma amplitude is uniformly distributed across all phases) and 1 (perfect phase–locked amplitude modulation, i.e., when the γ amplitude is highly concentrated in one phase).This calculation was repeated on a trial-by-trial basis for all 96 channels and for both conditions (*TNS_0_* and *TNS_1_*), resulting in a distribution of MI values per channel and condition. The direction of change was also recorded: +1 (*TNS_1_ > TNS_0_*), –1 (*TNS_0_ > TNS_1_*), or 0 (no significant difference). Results were displayed as 8×12 spatial heatmaps matching the physical layout of the electrode array, summarizing p-values, significance, and directional effects across all channels.

To observe the differences between *TNS_0_* and *TNS_1_* (in Figure 4), the MI was computed for all the trial duration (- 0.05 to 1 sec) and then to temporally dissociate these differences, the MI was calculated in 8 half-overlapping windows (in Figure 5) from TNS onset (0) to 1 sec (trial end).

### Statistical Models

To predict the post stimulation MI from the pre stimulation MI, we built the following Linear Mixed Effects (LME) model:

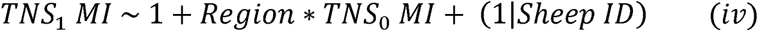

where *TNS_1_ MI* is the post-stimulation MI, ‘*Region*’ is a categorical variable indicating the cortical region (association vs. somatosensory cortex), and *TNS_0_ MI* is pre-stimulation (baseline) MI. The interaction term between ‘region’ and *TNS_0_ MI* tests whether the relationship between pre- and post-stimulation modulation differs across cortical regions. The model includes a random intercept for each individual sheep to account for variability across subjects.

To predict the MI across time steps we built another LME model as follows:

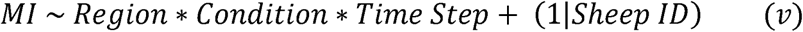

where *MI* is modulation index at TNS_0_ or TNS_1_, ‘*Condition’* (TNS_0_ or TNS_1_) is a categorical variable referring to indicating the cortical ‘*Region*’ (association vs. somatosensory cortex), and *Time Step* represents discrete temporal segments. The model includes all main effects and their interactions: *Region*, *Condition*, *TimeStep*, as well as all two-way and the three-way interaction terms among these factors. A random intercept for *SheepID* was included to account for inter-subject variability.

For each condition, the sum of two Gaussian functions was fit to the normalized modulation index across time using MATLAB’s ‘*gauss2*’ model (Curve Fitting Toolbox), with dynamic initialization of peak locations based on *‘findpeaks’*.

### Statistical tests

MI differences between *TNS_0_* and *TNS_1_*were assessed using the non-parametric Mann–Whitney U test (MATLAB ranksum). Radial plot statistics were computed using the MATLAB Circular Statistics Toolbox: theta phase locking was tested with the Rayleigh test, and condition differences with the Watson–Williams test. A permutation t-test (5000 permutations, Bonferroni corrected) assessed somatosensory vs. association channels, and a KS-test evaluated differences in temporal curves. Exact p-values are reported when > 0.001; otherwise, p < 0.001 is noted (Curran-Everett and Benos, 2007). A 95% confidence threshold (α = 0.05) was used to determine statistical significance.

## RESULTS

To investigate how hSCS modulates cortical oscillatory dynamics, we analyzed cross-frequency interactions between low-frequency (□: 4–8□Hz) and high-frequency (high γ: 70–150 Hz) activity in the somatosensory and association cortices of sheep. The overall -□γ locking across all included channels was quantified as the mean resultant length (MRL, i.e., it computes an average over all preferred angles across channels), providing a measure of the consistency of γ amplitude occurring at a preferred phase. We then computed MI, a measure of Phase-Amplitude Coupling Strength (PACS), on a trial-by-trial basis across two conditions: *TNS_0_* (pre-hSCS) and *TNS_1_* (post-hSCS). By comparing MI distributions across these conditions, we assessed whether hSCS altered the strength of PAC, a mechanism implicated in sensory integration and pain modulation (Wang et al., 2011; Taesler and Rose, 2016).

### □-**γ** Phase Locking and Its Strength

To evaluate the overall modulation of theta-gamma locking across channels, we first calculated the □-γ locking for each channel and then computed the mean resultant lengths (MRL, indicated by the magenta solid line) across channels at *TNS_0_* and *TNS_1_* conditions. MRL quantifies the strength or consistency of □-γ phase-locking across channels, while the preferred phase angle indicates the phase angle at which γ amplitude is maximal. In Sheep1 (**Fig. 2A**), the MRL slightly decreased from 0.51 to 0.48 post-stimulation, indicating that the strength of -γ locking remained largely unchanged. However, significant locking was observed in both conditions (Rayleigh test; TNS_0_: *p* < 0.001; TNS_1_: *p* < 0.001). No significant difference in the mean preferred phase was found between *TNS_0_* and *TNS_1_* (Watson-Williams test, *p* = 0.83), suggesting hSCS did not alter cortical phase preference. Similar results were obtained for other sheep. In Sheep2 (**Fig. 2B**), the MRL slightly decreased from 0.39 to 0.31 after stimulation with consistent locking (Rayleigh test; TNS_0_: *p* < 0.001; TNS_1_: *p* < 0.001) and no significant change in preferred phase post-stimulation (Watson-Williams test, *p* = 0.44). In Sheep 3 (**Fig. 2C**) and Sheep 4 (**Fig. 2D**), the MRL increased from 0.40 to 0.52 and from 0.59 to 0.63 post-stimulation, respectively, indicating stronger -γ locking. Both sheep exhibited robust phase-locking across conditions (Sheep3 — TNS_0_: *p* < 0.001; TNS_1_: *p* < 0.001; Sheep 4 — TNS_0_: *p* < 0.001; TNS_1_: *p* < 0.001) with no shift in average preferred phase (Sheep3: *p* = 0.91; Sheep 4: *p* = 0.78), indicating preserved phase preference following hSCS. An example -γ phase-amplitude plots for Sheep 3 is shown in **Supplementary Figure 1.**

**Figure 2:**
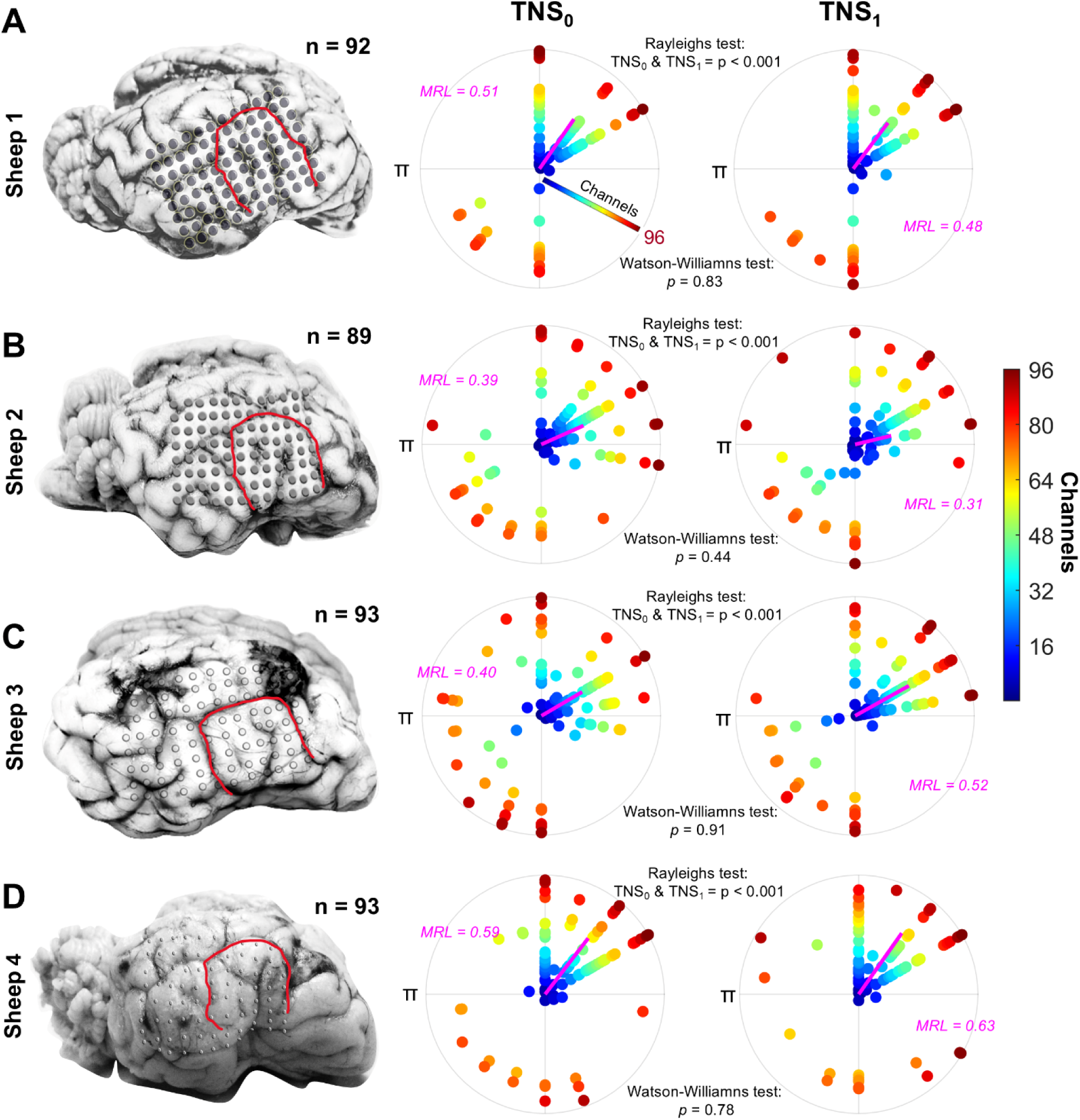
Preferred phase of γ modulation ( -γ locking) pre- (TNS_0_) and post-stimulation (TNS_1_) across sheep. **(A)** Left: Anatomical location of the ECoG electrodes in Sheep 1. The red line indicates the border between Somatosensory (inside) and Association cortices. Middle: Polar scatter plots show channel-wise preferred phases for TNS_0_, with mean resultant length (MRL) indicated by magenta lines. Right: Polar scatter plots (with MRL) show channel-wise preferred phases for TNS_1_. (B-D) Same as A but for Sheep 2 **(B)**, Sheep 3 **(C)** and Sheep 4 **(D)**. Across all animals, MRLs remained comparable, reflecting consistent theta-gamma locking. No significant shift in preferred theta phase was observed (Watson-Williams test, p > 0.05), suggesting hSCS preserved cortical phase preference. The color scale represents channel indices, ranging from 1 (blue) to 96 (dark red). For clarity the color scale is also embedded in A (middle). ‘n’ indicates number of channels included for analysis.

Overall, hSCS modulated the strength -γ locking in some sheep but did not significantly alter the average preferred phase across the cortical surface.

### □-**γ** Locking: Somatosensory vs. Association Cortices

We then proceeded with comparing the □-γ locking for the somatosensory *vs.* association cortex across all sheep **(Fig. 3).** In the association cortex **(Fig. 3A-D**), robust and consistent -γ locking was observed in all sheep. In Sheep 1 **(Fig. 3A**), MRLs remained high for *TNS_0_*(0.79) and *TNS_1_* (0.73), with highly significant □-γ locking (Rayleigh results, *p* < 0.001) and no significant difference in the preferred phase (Watson-Williams test, *p = 0.56*). Sheep 2 **(Fig. 3B**) showed moderate MRLs (TNS_0_ = 0.59, TNS_1_ = 0.44) and significant □-γ locking (Rayleigh test, p < 0.001) with no significant phase shift (Watson-Williams test, p = 0.23). In Sheep 3 **(Fig. 3C**), MRL decreased slightly from 0.55 to 0.45, but both conditions showed strong -γ locking (Rayleigh test, p < 0.001), and no phase shift (Watson-Williams test, p = 0.85). Sheep 4 **(Fig. 3D**) exhibited strong □-γ locking in both states (MRL = 0.7574 and 0.7792; Rayleigh p < 0.001), again without significant change in preferred phase (Watson-Williams test, p = 0.49).

**Figure 3.**
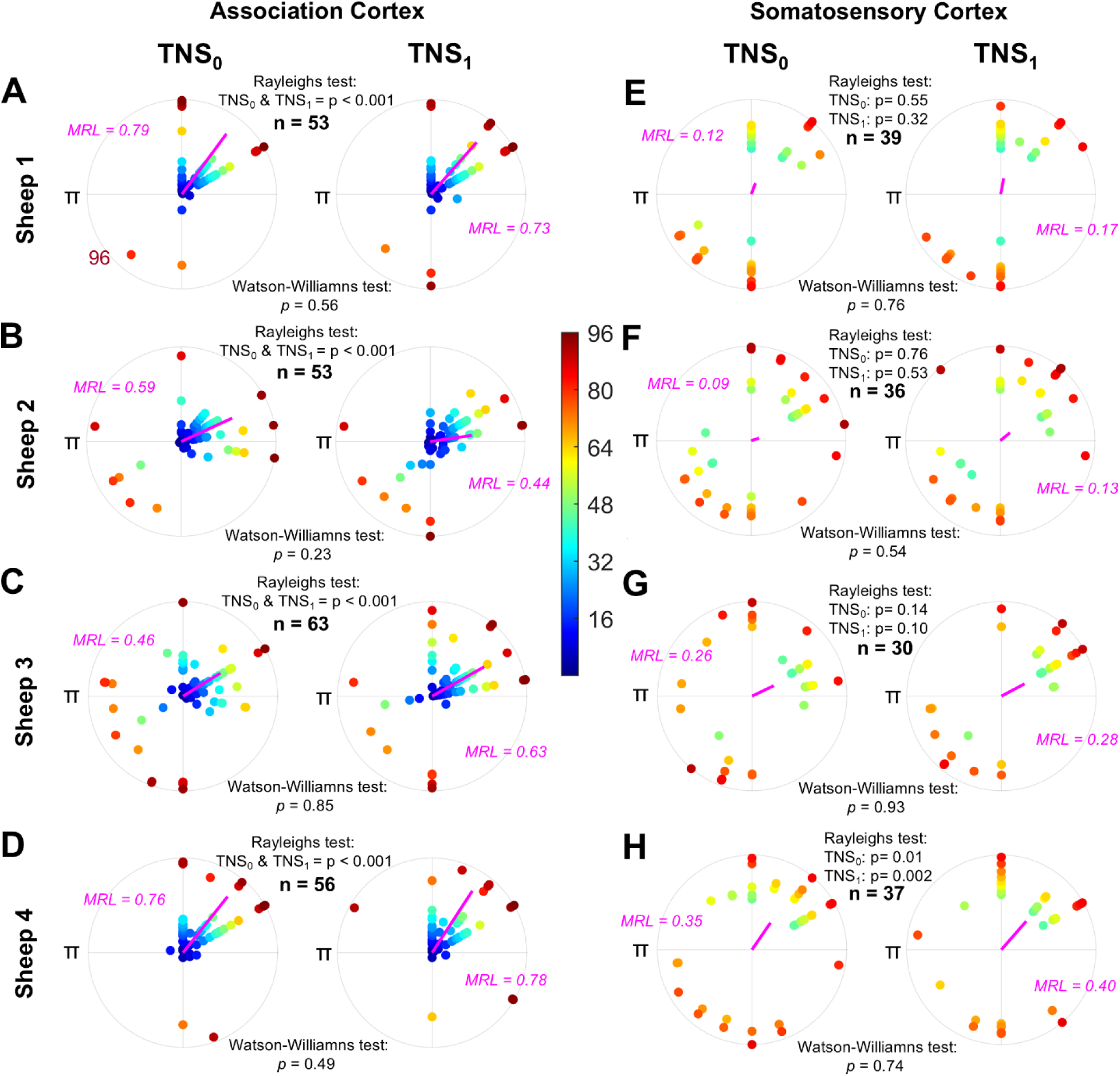
Comparison of -γ locking in association vs. somatosensory cortex across all sheep. **(A–D)** In the association cortex, strong and consistent -γ locking was observed across all sheep, with high mean resultant lengths (MRLs), significant phase locking (Rayleigh p < 0.001), and no significant shift in preferred phase following stimulation (Watson-Williams p > 0.05). **(E–H)** In contrast, the somatosensory cortex showed weak and inconsistent -γ locking, with low MRLs, mostly non-significant phase locking, and no significant phase shift between TNS_0_ and TNS_1_. The color scale represents channel indices, ranging from 1 (blue) to 96 (dark red).

**Figure 4.**
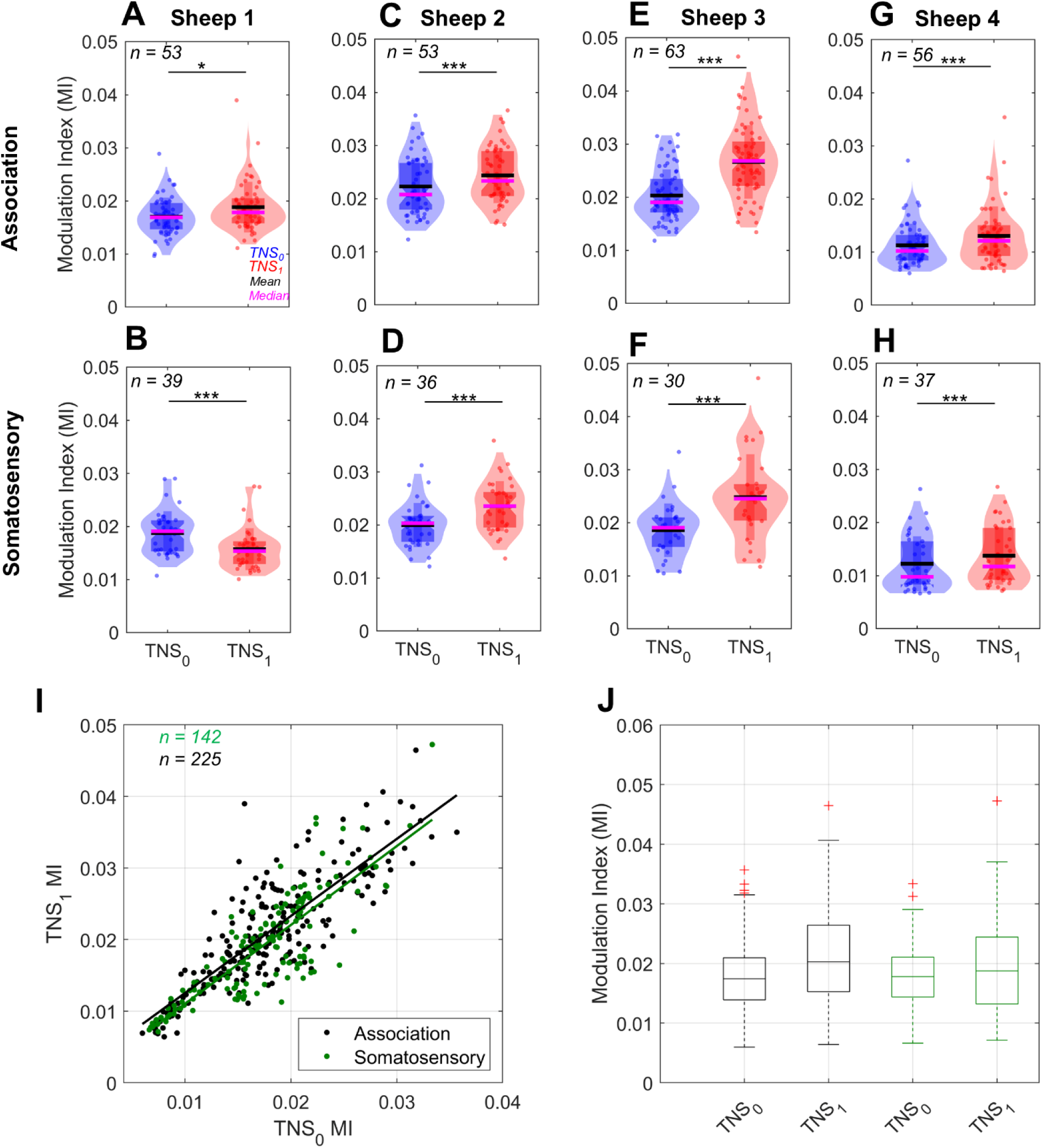
-γ coupling strength using Modulation index (MI) (A–B) In Sheep 1. (**A**), MI significantly increased in the association cortex from TNS0 to TNS1 but decreased in the somatosensory cortex (**B**). (C–H) In Sheep 2–4, MI increased post-stimulation in both the association (**C, E, G**) and somatosensory (**D, F, H**) cortices, with all comparisons showing statistical significance (permutation t-tests, p < 0.05). (I–J) Linear mixed-effects modeling revealed that TNS0 MI was significantly lower in the somatosensory cortex compared with the association cortex (p = 0.013), while TNS0 MI strongly predicted TNS1 MI (p < 0.001). A trend toward interaction between cortical region and TNS0 MI was observed, though not statistically significant (p = 0.08). Random intercepts by subject accounted for individual variability. Overall, these findings suggest that hSCS enhances phase-amplitude coupling, particularly in the association cortex.

**Figure 5.**
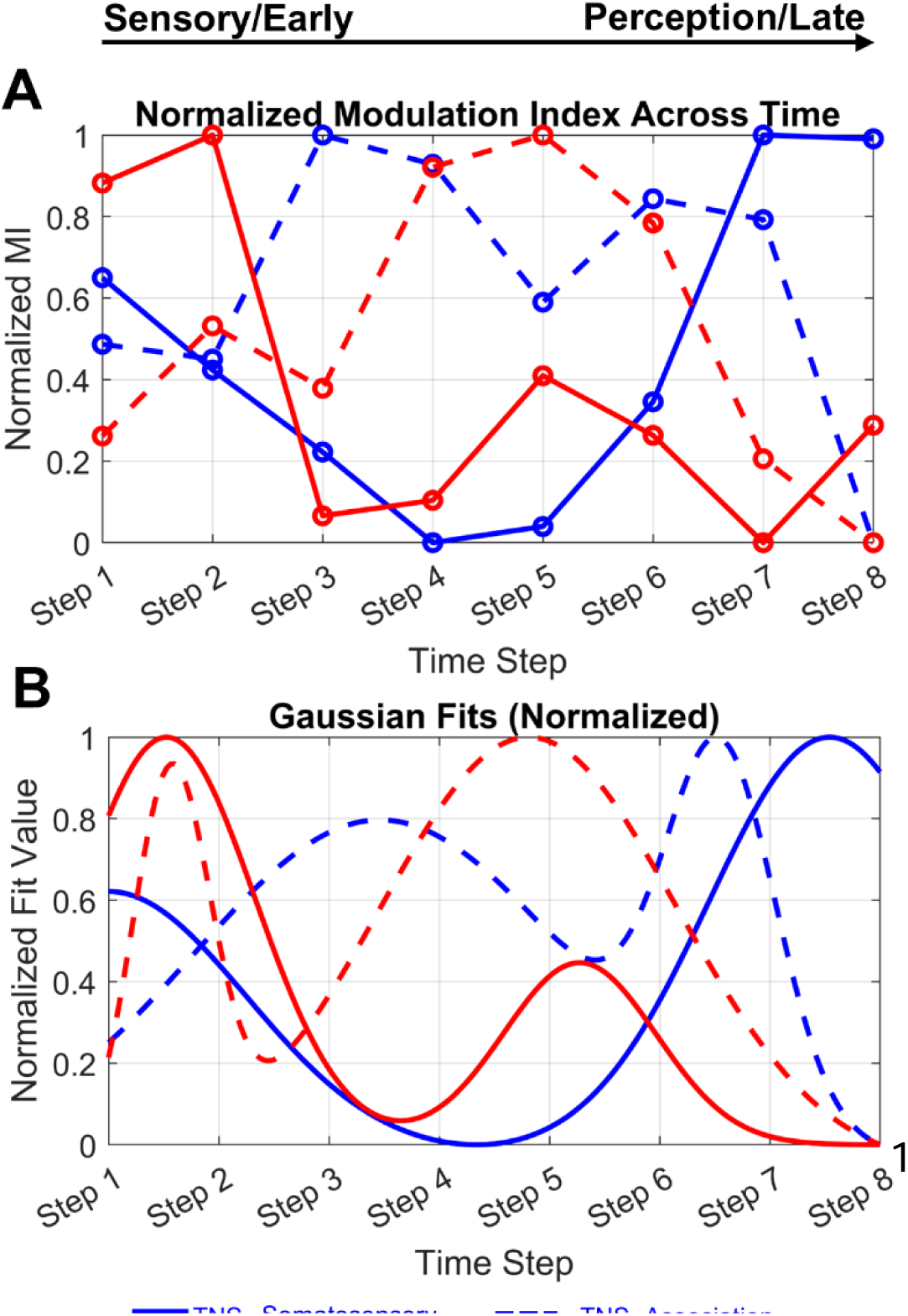
Temporal changes in modulation index (MI) after hSCS stimulation. **(B)** Normalized data highlights differences in peak MI across conditions and cortices. **(C)** Gaussian fits reveal a significant forward shift in peak MI from TNS_0_ to TNS_1_ in both cortices (KS test; Association: p = 0.0004, Somatosensory: p = 0.0018), indicating a reciprocal modulation of cortical activity.

Contrastingly, in the somatosensory cortex, □-γ locking was consistently weak across all sheep **(Fig. 3E-H)**. In Sheep 1 **(Fig. 3E**), the MRL was low both *TNS_0_* (0.13) and *TNS_1_* (0.17), with non-significant locking (Rayleigh test, TNS_0_: p = 0.547 and TNS_1_: p = 0.32) and no difference in preferred phase (Watson-Williams test, p = 0.76). Similarly, Sheep 2 **(Fig. 3F**) showed low MRLs (TNS_0_ = 0.09, TNS_1_ = 0.13), non-significant locking (Rayleigh tests, p = 0.76 and 0.53, respectively), and no significant phase shift (Watson-Williams test, p = 0.54). In Sheep 3 **(Fig. 3G**), both conditions showed minimal locking (MRL = 0.26 and 0.28; Rayleigh p = 0.79 and 0.79 respectively) and no significant phase difference (Watson-Williams p = 0.93). Sheep 4 **(Fig. 3H**) exhibited slightly stronger MRLs and locking (TNS_0_ = 0.35, TNS_1=_ 0.40; Rayleigh p = 0.01 and 0.002, respectively), yet again with no significant phase shift (p = 0.74).

Together, these findings demonstrate that stimulation did not significantly alter the preferred phase of γ activity in either cortical region. Importantly, a consistent dissociation emerged: -γ locking was robust and reliable in the association cortex, whereas it remained weak and often non-significant in the somatosensory cortex.

### Modulation index: Somatosensory vs. Association cortex

Having established that the preferred phase angle does not significantly differ between *TNS_0_ and TNS_1_* for both cortices, we next examined changes in the modulation index (MI), a measure of □-γ coupling strength. Sheep 1 **(Fig. 4A)** showed differential modulation of -γ coupling between cortical areas. In the association cortex, MI significantly (permutation t-test, p = 0.017, t = -2.44) increased from *TNS_0_*(mean ± SEM = 0.0170 ± 0.0005, median = 0.0169) to *TNS_1_*(mean ± SEM = 0.0188 ± 0.0007, median = 0.0178). Conversely, in the somatosensory cortex **(Fig. 4B),** MI decreased (permutation t-test, p < 0.001, t = 6.84) from TNS_0_ (mean ± SEM = 0.0188 ± 0.0007, median = 0.0191) to TNS_1_ (mean ± SEM = 0.0157 ± 0.0006, median = 0.0154). Channel-by channel results for all sheep are shown in **supplementary Figure 2.**

For all other sheep **(Fig. 4C-H)** we observed a similar trend for both cortices. In the association cortex **(Fig 4C, E, G),** MI significantly increased (permutation t-test, p < 0.001; Sheep 2: t = - 9.06. Sheep 3: t = - 11.70, Sheep 4: - 6.40) from *TNS_0_* (Sheep 2: mean ± SEM = 0.0223 ± 0.0007, median = 0.0208; Sheep3: mean ± SEM = 0.0203 ± 0.0006, median = 0.0190; Sheep 4: mean ± SEM = 0.0113 ± 0.0005, median = 0.0102) to *TNS_1_* (Sheep 2: mean ± SEM = 0.0244 ± 0.0007, median = 0.0233; Sheep3: mean ± SEM = 0.0266 ± 0.0009, median = 0.0268; Sheep 4:mean ± SEM = 0.0130 ± 0.0007, median = 0.0121). Similarly, the somatosensory cortex **(Fig 4D, F, H)** showed a significant increase (permutation t-test, p < 0.001; Sheep 2: t = - 18.84. Sheep 3: t = - 9.38, Sheep 4: - 8.55) in MI from *TNS_0_* (Sheep 2: mean ± SEM = 0.0200 ± 0.0007, median = 0.0203; Sheep3: mean ± SEM = 0.0186 ± 0.0009, median = 0.0191; Sheep4: mean ± SEM = 0.0122 ± 0.0009, median = 0.0098) to *TNS_1_* (Sheep 2: mean ± SEM = 0.0235 ± 0.0008, median = 0.0236; Sheep 3: mean ± SEM = 0.0248 ± 0.0015, median = 0.0245; Sheep 4: mean ± SEM = 0.0137 ± 0.0009, median = 0.0117).

We then used a linear mixed-effects model to investigate how baseline MI (TNS_0_ MI) and cortical region (association vs. somatosensory cortex) predict post-stimulation MI (TNS_1_ MI), while accounting for inter-animal variability. Overall, the model exhibited a strong fit (AIC = −3097.7, BIC = −3074.2, log-likelihood = 1554.8, and deviance = −3109.7) **(Figure 4)**.

The linear mixed-effects model revealed differences between the association and somatosensory cortices, with the somatosensory cortex exhibiting a significantly lower TNS_1_MI compared with the association cortex (fixed effect: β = −0.0030, *p* = 0.013). The TNS_0_ MI was a strong positive predictor of TNS_1_MI across both cortical areas (fixed effect: β = 0.947, *p* < 0.001), indicating robust consistency in coupling strength following stimulation. The interaction between cortical area and TNS_0_MI showed a trend toward significance (β = 0.11, *p* = 0.08), suggesting a possible difference in the slope of the TNS_0_–TNS_1_ MI relationship between the somatosensory and association cortices. However, the effect did not reach statistical significance, indicating that the regression lines are largely parallel, with only a slight and non-significant difference in slope across regions.

The model included random intercepts for each sheep to account for individual baseline differences in post-stimulation modulation index (TNS_1_MI). The estimated variance of these random intercepts was 0.00237 (95% CI [0.00116, 0.00482]), indicating modest variability between animals. The residual variance, representing within-subject variability (e.g., across cortical regions and conditions), was larger at 0.00343 (95% CI [0.00319, 0.00368]). These results suggest that although baseline differences exist between sheep, most of the variability in TNS_1_ MI arises from within-subject factors rather than between-subject differences. Thus, the model effectively accounts for biological variability across individuals while highlighting the influence of cortical area and stimulation condition.

### Temporal evolution of modulation index: Somatosensory vs. Association cortex

Until now, we have established that -γ phase-amplitude locking remains largely stable across cortical regions, with some subtle regional differences, while the MI was significantly modulated following hSCS. Specifically, this trend was similar across three sheep except for Sheep 1. What is the effect in the temporal domain, i.e., how does the effect develop in time? To explore how this effect evolves over time, we divide the trial data into 8 half-overlapping steps

Figure 5 illustrates the temporal dynamics of MI across cortical areas and stimulation conditions. Figure 5A shows normalized values (0–1), highlighting distinct peaks in MI between cortices. Two temporal features emerge: the somatosensory cortex exhibits a rapid MI increase immediately after hSCS (notably at time steps 1–2), while the association cortex shows a delayed, gradual enhancement (time steps 4–6). This lagged response likely reflects region-specific differences in synaptic or network-level recruitment. Figure 5B presents Gaussian fits to the normalized data, further emphasizing these inter-regional temporal distinctions from TNS0 to TNS1. Individual and pooled average MI temporal trajectories for sheep are shown in **Supplementary** Figure 3A-D **and** Figure 3E, respectively.

In the somatosensory cortex, the higher peak observed at time steps 6–8 at TNS_0_ (alongside a smaller peak at steps 1 and 2, corresponding to the evoked response period) shifts to a pronounced peak at time steps 1 and 2 at TNS_1_. A similar pattern is seen in the association cortex, with a more pronounced peak at steps 1 and 2 at TNS_1_. Notably, these transitions are significant between conditions in both areas (two-sample, two-sided KS test; Association: p < 0.001, Somatosensory: p = 0.0018), suggesting a robust reciprocal shift of peaks from TNS_0_ to TNS_1_.

To further investigate the temporal dynamics of modulation index (MI) changes across cortical regions and stimulation conditions, we fitted a linear mixed-effects model with MI as the dependent variable. The model included fixed effects for ‘*Region*’ (Association vs. Somatosensory), ‘*Condition*’ (TNS_0_ vs TNS_1_), ‘*Time Step’* (categorical), and their interactions, as well as a random intercept for ‘*Sheep*’ to account for inter-subject variability. The model showed a good fit to the data (AIC = –1116.5, Log Likelihood = 592.25) with 128 observations.

The intercept corresponds to the baseline MI at the Association TNS_0_ condition at the first Time Step (estimate = 0.0504, p < 0.0001). A significant main effect of *‘Region’* indicated that the somatosensory cortex had a higher baseline MI than the association cortex during TNS_0_ (estimate = 0.00365, p = 0.021). The main effect of ‘*Condition***’** (TNS_0_ vs TNS_1_) was not significant overall (estimate = 0.00036, p = 0.81), suggesting no uniform shift in MI across all regions following stimulation.

Although the main effect of ‘*Time Step’* alone did not reach statistical significance (p > 0.05), the interaction terms revealed nuanced temporal dynamics that differ depending on both cortical region and stimulation condition. This suggests that MI changes over time are not simply linear or uniform across all experimental conditions but rather depend on the specific combination of region and condition. This complexity reflects the intricate nature of cortical responses to stimulation, where distinct mechanisms may govern MI modulation in the association *vs.* somatosensory cortices and during TNS_0_ versus TNS_1_ stimulation periods. The lack of widespread significant interactions also points to the potential influence of inter-subject variability (random intercept standard deviation ≈ 0.0047) and the need for further investigation with larger sample sizes or complementary analyses to fully characterize these temporal patterns.

These results further corroborate MI dynamics illustrated in Figure 5, where the somatosensory cortex exhibited a higher baseline MI and a rapid increase in MI post-stimulation followed by stabilization, whereas the association cortex showed lower baseline MI and more gradual temporal modulation. Collectively, the linear mixed-effects model supports distinct and reciprocal temporal modulation of MI across cortical areas and stimulation conditions following hSCS.

## DISCUSSION

We investigated the effect of hSCS on the -γ phase-amplitude locking and MI in the sheep association and somatosensory cortices. We show that hSCS: 1) did not change the overall - locking in both cortices, however 2) the MI was significantly modulated in both cortices (mostly increased). 3) Temporally, both cortices showed a reciprocal shifting of the MI from pre- to post-hSCS stimulation, and 4) finally a linear mixed model confirmed the complementary (reciprocal shifting of the response peaks in the temporal domain) cortical dynamics and distinct temporal evolution of MI over time.

### Methodological Considerations

Our experiments were conducted on anesthetized animals; therefore, no attentional parameters or behavioral readouts were involved. However, this preparation allowed for stable recordings through precise stimulation. It is important to note that different oscillatory frequencies can be observed under anesthesia (Gray and Singer, 1989; Lisman and Jensen, 2013; Bharmauria et al., 2015). Moreover, no sensitization / desensitization processes occurred during the stimulation conditions, as previously reported (Bharmauria et al., 2025). While our stimulation protocols were designed and optimized (Flouty et al., 2012; Kapural et al., 2015; Arle et al., 2016; Sdrulla et al., 2018; Rogers et al., 2022) to target tibial and spinal afferents, both TNS (Jørum and Shyu, 1988; Paquette and Yoo, 2019) and hSCS likely recruited a mixed population of sensory fibers (Head et al., 2019; Abraham et al., 2021; Rogers et al., 2022). Prior studies suggest that even low-frequency peripheral nerve stimulation can activate both nociceptive and mechanoreceptive afferents (Jørum and Shyu, 1988; Paquette and Yoo, 2019).

### □-**γ** Coupling, Modulation index and E/I balance: Functional implications

We have shown in prior studies that high γ oscillations rise with increasing peripheral stimulus intensity (Flouty et al., 2013), establishing an indirect link between physiology and experience: stronger shocks drive larger evoked responses and higher γ power, serving as biomarkers that correspond to, but are distinct from, the subjective perception of pain intensity (Wang et al., 2011; Lisman and Jensen, 2013; Taesler and Rose, 2016). Consistent with earlier findings, hSCS robustly suppressed high-frequency oscillations (i.e., strengthening of the inhibition) across both somatosensory and association cortices, reflecting a shift in the excitation/inhibition (E/I) balance (Kann, 2016; Ploner et al., 2017; Tan et al., 2019; Parker et al., 2020; Tan and Kuner, 2021; Ma and Khadra, 2024) likely mediated by altered local inhibitory circuits and neuronal synchronization (Ratnadurai-Giridharan et al., 2015; Kann, 2016). The observation that hSCS attenuates high-frequency oscillations and cortical hyperexcitability (Oshiro et al., 2009; Kapural et al., 2015; Caylor et al., 2019), in parallel with reduction in pain perception supports high γ activity as an indirect biomarker of pain and, more broadly, sensory processing.

Importantly, our detailed analyses of -γ cross-frequency coupling revealed that hSCS significantly modulated the coupling strength (MI) (Figure 4) without changing the preferred phase or -γ locking (Figures 2-3). Across subjects and cortical areas, -γ locking remained robust and consistent in the association cortex, with generally weaker locking in the somatosensory cortex, suggesting that hSCS selectively influences the gain or amplitude modulation of γ oscillations relative to phase, without disrupting the temporal coordination of cortical oscillations (Canolty and Knight, 2010; Tort et al., 2010; Lisman and Jensen, 2013; Aru et al., 2015; Fries, 2015).

Despite stable □-γ phase locking, hSCS increased the MI, enhancing cross-frequency interaction strength while preserving timing, suggesting sharper –γ coupling and potentially improved inhibitory control or gain modulation. (Canolty and Knight, 2010; Akam and Kullmann, 2014; Voytek and Knight, 2015). This sharpening could manifest as greater suppression of γ amplitude at non-preferred phases (i.e., the flanking bins), indicating more selective temporal gating of cortical excitability. Our results resemble neuroplastic processes, where stimulation or experience can sharpen or broaden neuronal tuning (Isaacson and Scanziani, 2011; Froemke, 2015; Bharmauria et al., 2019, 2022). Such modulation adjusts sensory encoding via precise inhibitory control over preferred and non-preferred inputs.

Briefly, hSCS refines the precision of cross-frequency interactions without altering their timing, optimizing sensory processing while preserving core oscillatory dynamics. This selective amplitude tuning via inhibitory gating may enhance cortical excitability and information flow between association and somatosensory regions, supporting therapeutic efficacy (Sohal et al., 2009; Isaacson and Scanziani, 2011; Akam and Kullmann, 2014; Voytek and Knight, 2015). hSCS preserves the basic cross-frequency architecture of sensory processing, with low-frequency phases still organizing high-frequency activity. High γ power remains a robust correlate of sensory processing, scaling with the intensity of peripheral stimulation, while locking to the preferred phase conveys qualitative sensory information (Flouty et al., 2013; Lisman and Jensen, 2013; Fries, 2015).

### Possible neural mechanisms: Association vs. Somatosensory cortex

Distinct patterns observed between association and somatosensory cortices highlight their specialized roles. In the association cortex, strong and consistent -γ coupling suggests that low-frequency rhythms temporally organize high-frequency bursts (Canolty and Knight, 2010; Lisman and Jensen, 2013). This phase-amplitude interaction likely supports selective attention and multimodal integration (Buzsáki, 2006; Oshiro et al., 2009; Zhang et al., 2014; Tan and Kuner, 2021) by aligning excitatory inputs to optimal phases of slower oscillations (Lisman and Jensen, 2013). In contrast, somatosensory cortex exhibited more variable phases, indicating locally driven, stimulus-dependent modulation (Canolty et al., 2006; Sirota et al., 2008; Zagha et al., 2013; Darainy et al., 2023; Ebrahimi and Ostry, 2024). These differences align with their hierarchical roles and prior findings that low-frequency rhythms more strongly organize high-frequency activity in association than in primary sensory regions (Engel et al., 2001; Gilbert and Sigman, 2007). This dual role likely increases variability in phase–amplitude relationships, allowing network states to update with sensory inputs while remaining aligned with contextual signals (Bastos et al., 2012). These results are consistent with prior findings that low-frequency rhythms more strongly organize high-frequency activity in the association cortex than in the somatosensory cortex (Bharmauria et al., 2025) and with fMRI reports of enhanced BOLD variability and altered connectivity in somatosensory regions under pain (Kim et al., 2020; Lim et al., 2021). These differences point to a diminished role for slow oscillations in orchestrating activity in primary sensory areas, consistent with their stimulus-locked, bottom-up processing style (Gilbert and Sigman, 2007; Fries, 2015; Keitel and Gross, 2016).

Interestingly, in the temporal domain, hSCS elicited a rapid but transient increase in MI in the somatosensory cortex at early time steps (1 and 2), followed by a reduction in amplitude and a shift in the second, delayed MI peak. In contrast, the association cortex showed not only a transient early increase and sharpening of the initial peak (time steps 1 and 2), but also a broadening and earlier shift of the second peak from time steps 6 and 7 to 4 and 5. Overall, these reciprocal shifts in MI peak timing between the two regions suggest a sequential and functionally differentiated engagement of somatosensory and association circuits following hSCS. This dynamic organization likely reflects a recalibration of sensory and perceptual processing, as earlier components of EPs are typically associated with initial sensory detection, whereas the later components are more linked to the perceptual awareness of a stimulus (Näätänen and Picton, 1987; Dehaene and Changeux, 2011; Bermudez-Contreras et al., 2023). It appears that, in our case, MI enhancement in the somatosensory area is representative of resetting for proper stimulus retrieval, with concurrent decrease in the perception (the second peak decrease and shift). Accordingly, the association cortex sensory processing also shifts and improves along with a more organized perception of the stimulus.

Together, these results extend our previous work by illustrating that hSCS modulates cortical oscillations at multiple levels: suppressing excessive high-frequency activity, adjusting cross-frequency coupling strength while preserving phase-locking stability with a dynamic retuning of the sensory/perception loop in the temporal domain (Näätänen and Picton, 1987; Canolty and Knight, 2010; Dehaene and Changeux, 2011; De Ridder et al., 2013; Voytek and Knight, 2015). This multifaceted modulation likely contributes to the normalization of cortical excitability and sensory perception underlying the analgesic effects of hSCS.

### Limitations and future directions

Several limitations warrant consideration. First, the use of an anesthetized ovine model may not fully capture the complexity of awake cortical dynamics. However, the use of well-established anesthesia protocols allowed us to achieve stable, high quality multichannel cortical recordings over long experimental durations, a methodological trade-off that enabled precise quantification of cross-frequency interactions across distributed networks. To bridge this gap, future investigations should leverage chronic, awake models that allow for real-time behavioral correlations. For example, a simplified translation of this approach in humans (e.g., electroencephalography combined with TNS and hSCS) could provide an important next step for testing this mechanism. Future studies could investigate state-dependent differences by comparing network behavior under anesthesia versus awake conditions, as observed in DBS patient recordings (Shirvalkar et al., 2020; Thibes et al., 2025). Additionally, linking oscillatory changes to pain relief in a hypothesis-driven manner, and exploring long-term effects of hSCS while optimizing stimulation parameters to selectively target afferent pathways, will be crucial for enhancing clinical relevance and translational impact. Integrating molecular markers and optogenetic techniques could also offer mechanistic insight into how hSCS shapes cortical oscillatory activity and alters pain perception (Gao and Ji, 2009; Smits et al., 2009; Liu et al., 2016; Fujimori et al., 2022). Second, the sample size (N=4) reflects ethical and technical constraints, but permutation tests and mixed-effects models accounted for variability, and consistent patterns across animals support the robustness of our findings. Notably, the current analysis focused on intra-electrode cross-frequency coupling; however, future studies could expand this scope by investigating inter-electrode coupling to uncover spatial coordination across cortical sites (Tort et al., 2008, 2010; Hyafil et al., 2015). In a similar vein, adopting a network-analysis framework may help disentangle the functional organization of these two complementary brain regions, revealing potential hubs and modules that mediate the effects of hSCS (Sporns and Betzel, 2016; Ghaderi et al., 2020).

## Conclusion

In conclusion, this study demonstrates the dynamic and multidimensional effects of hSCS on cortical oscillatory dynamics in the anesthetized ovine brain. These findings shed light on how pain may be processed and modulated at the cortical level through both local and supraspinal mechanisms. Overall, the observed reorganization likely reflects a recalibration of the sensory-perceptual processing loop, with implications for how stimuli are sensed, integrated, and interpreted in the cortex. By preserving rhythmic phase structure while modulating phase-specific amplitude dynamics, hSCS emerges as a promising neuromodulatory approach capable of restoring cortical balance and potentially alleviating chronic pain. By bridging local physiology with systems-level organization, this work lays critical groundwork for future translational studies in awake and behaving models, and ultimately for the development of biomarker-guided, adaptive neuromodulation strategies in humans.

## Author contributions

VB conducted data analysis with contributions from OF, NS and AGh. YB, NS, AGh, KJ, HO, HK, YYB, AS, MAH, CS, BD, and AML contributed to data interpretation, writing and editing. VB and OF drafted the manuscript, with revisions and inputs from all authors. HO, HK, MAH, OF conceptualized the study and did the experiments. MAH and OF are cosenior authors.

## Disclosures

This study was supported by a grant from the University of Iowa to MH III and the University of South Florida seed grant to OF. No conflict of interest for any of the authors. Study registration This study protocol has been registered with University of Iowa (IACUC# 0902039).

## Data availability

All data can be found in the main text or supplementary materials. Additional materials related to this project are available upon request.

## Notes

### Competing Interest Statement

The authors have declared no competing interest.

## References

1. Abraham, M. E., Gold, J., Dondapati, A., Sheaffer, K., Gendreau, J. L., and Mammis, A. (2021). High Frequency 10 kHz Spinal Cord Stimulation as a First Line Programming Option for Patients With Chronic Pain: A Retrospective Study and Review of the Current Evidence. Cureus 13, e17220. doi: 10.7759/cureus.17220

2. Akam, T., and Kullmann, D. M. (2014). Oscillatory multiplexing of population codes for selective communication in the mammalian brain. Nat Rev Neurosci 15, 111–122. doi: 10.1038/nrn3668

3. Arle, J. E., Mei, L., Carlson, K. W., and Shils, J. L. (2016). High-Frequency Stimulation of Dorsal Column Axons: Potential Underlying Mechanism of Paresthesia-Free Neuropathic Pain Relief. Neuromodulation 19, 385–397. doi: 10.1111/ner.12436

4. Aru, J., Aru, J., Priesemann, V., Wibral, M., Lana, L., Pipa, G., et al. (2015). Untangling cross-frequency coupling in neuroscience. Curr Opin Neurobiol 31, 51–61. doi: 10.1016/j.conb.2014.08.002

5. Bastos, A. M., Usrey, W. M., Adams, R. A., Mangun, G. R., Fries, P., and Friston, K. J. (2012). Canonical microcircuits for predictive coding. Neuron 76, 695–711. doi: 10.1016/j.neuron.2012.10.038

6. Bermudez-Contreras, E., Schjetnan, A. G.-P., Luczak, A., and Mohajerani, M. H. (2023). Sensory experience selectively reorganizes the late component of evoked responses. Cereb Cortex 33, 2626–2640. doi: 10.1093/cercor/bhac231

7. Bharmauria, V., Bachatene, L., Cattan, S., Chanauria, N., Rouat, J., and Molotchnikoff, S. (2015). Stimulus-dependent augmented gamma oscillatory activity between the functionally connected cortical neurons in the primary visual cortex. European Journal of Neuroscience 41, 1587–1596. doi: 10.1111/ejn.12912

8. Bharmauria, V., Bachatene, L., and Molotchnikoff, S. (2019). The speed of neuronal adaptation: A perspective through the visual cortex. European Journal of Neuroscience 49. doi: 10.1111/ejn.14393

9. Bharmauria, V., Ouelhazi, A., Lussiez, R., and Molotchnikoff, S. (2022). Adaptation-induced plasticity in the sensory cortex. J Neurophysiol 128, 946–962. doi: 10.1152/jn.00114.2022

10. Bharmauria, V., Oya, H., Bezchlibnyk, Y., Shaheen, N., Ghaderi, A., Johari, K., et al. (2025). Neurophysiological effects of high-frequency spinal cord stimulation on cortico-sensory areas in large ovine animal model. J Pain 34, 105493. doi: 10.1016/j.jpain.2025.105493

11. Buzsáki, G. (2006). Rhythms of the Brain. Oxford University Press. Available at: https://global.oup.com/academic/product/rhythms-of-the-brain-9780199828234?cc=ca&lang=en&

12. Canolty, R. T., Edwards, E., Dalal, S. S., Soltani, M., Nagarajan, S. S., Kirsch, H. E., et al. (2006). High gamma power is phase-locked to theta oscillations in human neocortex. Science 313, 1626–1628. doi: 10.1126/science.1128115

13. Canolty, R. T., and Knight, R. T. (2010). The functional role of cross-frequency coupling. Trends Cogn Sci 14, 506–515. doi: 10.1016/j.tics.2010.09.001

14. Caylor, J., Reddy, R., Yin, S., Cui, C., Huang, M., Huang, C., et al. (2019). Spinal cord stimulation in chronic pain: evidence and theory for mechanisms of action. Bioelectron Med 5, 12. doi: 10.1186/s42234-019-0023-1

15. Chakravarthy, K., Nava, A., Christo, P. J., and Williams, K. (2016). Review of Recent Advances in Peripheral Nerve Stimulation (PNS). Curr Pain Headache Rep 20, 60. doi: 10.1007/s11916-016-0590-8

16. Curran-Everett, D., and Benos, D. J. (2007). Guidelines for reporting statistics in journals published by the American Physiological Society: the sequel. Adv Physiol Educ 31, 295–298. doi: 10.1152/advan.00022.2007

17. Darainy, M., Manning, T. F., and Ostry, D. J. (2023). Disruption of somatosensory cortex impairs motor learning and retention. J Neurophysiol 130, 1521–1528. doi: 10.1152/jn.00231.2023

18. De Ridder, D., Plazier, M., Kamerling, N., Menovsky, T., and Vanneste, S. (2013). Burst spinal cord stimulation for limb and back pain. World Neurosurg 80, 642–649.e1. doi: 10.1016/j.wneu.2013.01.040

19. Dehaene, S., and Changeux, J.-P. (2011). Experimental and theoretical approaches to conscious processing. Neuron 70, 200–227. doi: 10.1016/j.neuron.2011.03.018

20. Ebrahimi, S., and Ostry, D. J. (2024). The human somatosensory cortex contributes to the encoding of newly learned movements. Proc Natl Acad Sci U S A 121, e2316294121. doi: 10.1073/pnas.2316294121

21. Ehsanian, R., Wu, V., Grandhe, R., Valeriano, M., Petersen, T. R., Rivers, W. E., et al. (2024). A single-center real-world review of 10 kHz high-frequency spinal cord stimulation outcomes for treatment of chronic pain. Interv Pain Med 3, 100402. doi: 10.1016/j.inpm.2024.100402

22. Engel, A. K., Fries, P., and Singer, W. (2001). Dynamic predictions: oscillations and synchrony in top-down processing. Nat Rev Neurosci 2, 704–716. doi: 10.1038/35094565

23. Flouty, O. E., Oya, H., Kawasaki, H., Reddy, C. G., Fredericks, D. C., Gibson-Corley, K. N., et al. (2013). Intracranial somatosensory responses with direct spinal cord stimulation in anesthetized sheep. PLoS One 8, e56266. doi: 10.1371/journal.pone.0056266

24. Flouty, O., Oya, H., Kawasaki, H., Wilson, S., Reddy, C. G., Jeffery, N. D., et al. (2012). A new device concept for directly modulating spinal cord pathways: initial in vivo experimental results. Physiol Meas 33, 2003–2015. doi: 10.1088/0967-3334/33/12/2003

25. Flouty, O., Yamamoto, K., Germann, J., Harmsen, I. E., Jung, H. H., Cheyuo, C., et al. (2022). Idiopathic Parkinson’s disease and chronic pain in the era of deep brain stimulation: a systematic review and meta-analysis. J Neurosurg 137, 1821–1830. doi: 10.3171/2022.2.JNS212561

26. Fries, P. (2015). Rhythms for Cognition: Communication through Coherence. Neuron 88, 220–235. doi: 10.1016/j.neuron.2015.09.034

27. Froemke, R. C. (2015). Plasticity of cortical excitatory-inhibitory balance. Annu Rev Neurosci 38, 195–219. doi: 10.1146/annurev-neuro-071714-034002

28. Fujimori, K., Sekine, M., Watanabe, M., Tashima, R., Tozaki-Saitoh, H., and Tsuda, M. (2022). Chemogenetic silencing of spinal cord-projecting cortical neurons attenuates Aβ fiber-derived neuropathic allodynia in mice. Neurosci Res 181, 115–119. doi: 10.1016/j.neures.2022.05.001

29. Gao, Y.-J., and Ji, R.-R. (2009). c-Fos and pERK, which is a better marker for neuronal activation and central sensitization after noxious stimulation and tissue injury? Open Pain J 2, 11–17. doi: 10.2174/1876386300902010011

30. Ghaderi, A. H., Baltaretu, B. R., Andevari, M. N., Bharmauria, V., and Balci, F. (2020). Synchrony and Complexity in State-Related EEG Networks: An Application of Spectral Graph Theory. Neural Comput 32, 2422–2454. doi: 10.1162/neco_a_01327

31. Gilbert, C. D., and Sigman, M. (2007). Brain states: top-down influences in sensory processing. Neuron 54, 677–696. doi: 10.1016/j.neuron.2007.05.019

32. Gray, C. M., and Singer, W. (1989). Stimulus-specific neuronal oscillations in orientation columns of cat visual cortex. Proceedings of the National Academy of Sciences of the United States of America 86, 1698–1702.

33. Gross, J., Schnitzler, A., Timmermann, L., and Ploner, M. (2007). Gamma oscillations in human primary somatosensory cortex reflect pain perception. PLoS Biol 5, e133. doi: 10.1371/journal.pbio.0050133

34. Hagedorn, J. M., Romero, J., Thuc Ha, C., Bendel, M. A., and D’Souza, R. S. (2021). Paresthesia-Based Versus High-Frequency Spinal Cord Stimulation: A Retrospective, Real-World, Single-Center Comparison. Neuromodulation. doi: 10.1111/ner.13497

35. Head, J., Mazza, J., Sabourin, V., Turpin, J., Hoelscher, C., Wu, C., et al. (2019). Waves of Pain Relief: A Systematic Review of Clinical Trials in Spinal Cord Stimulation Waveforms for the Treatment of Chronic Neuropathic Low Back and Leg Pain. World Neurosurg 131, 264–274.e3. doi: 10.1016/j.wneu.2019.07.167

36. Huang, Q., Oya, H., Flouty, O. E., Reddy, C. G., Howard, M. A., Gillies, G. T., et al. (2014). Comparison of spinal cord stimulation profiles from intra- and extradural electrode arrangements by finite element modelling. Med Biol Eng Comput 52, 531–538. doi: 10.1007/s11517-014-1157-7

37. Hyafil, A., Giraud, A.-L., Fontolan, L., and Gutkin, B. (2015). Neural Cross-Frequency Coupling: Connecting Architectures, Mechanisms, and Functions. Trends Neurosci 38, 725–740. doi: 10.1016/j.tins.2015.09.001

38. Isaacson, J. S., and Scanziani, M. (2011). How inhibition shapes cortical activity. Neuron 72, 231–243. doi: 10.1016/j.neuron.2011.09.027

39. Johnson, Q., Borsheski, R. R., and Reeves-Viets, J. L. (2013). A Review of Management of Acute Pain. Mo Med 110, 74–79.

40. Jørum, E., and Shyu, B.-C. (1988). Analgesia by low-frequency nerve stimulation mediated by low-threshold afferents in rats. Pain 32, 357–366. doi: 10.1016/0304-3959(88)90048-6

41. Kaas, J. H. (1993). The functional organization of somatosensory cortex in primates. Ann Anat 175, 509–518. doi: 10.1016/s0940-9602(11)80212-8

42. Kaas, J. H. (2004). Evolution of somatosensory and motor cortex in primates. Anat Rec A Discov Mol Cell Evol Biol 281, 1148–1156. doi: 10.1002/ar.a.20120

43. Kann, O. (2016). The interneuron energy hypothesis: Implications for brain disease. Neurobiol Dis 90, 75–85. doi: 10.1016/j.nbd.2015.08.005

44. Kapural, L., Yu, C., Doust, M. W., Gliner, B. E., Vallejo, R., Sitzman, B. T., et al. (2015). Novel 10-kHz High-frequency Therapy (HF10 Therapy) Is Superior to Traditional Low-frequency Spinal Cord Stimulation for the Treatment of Chronic Back and Leg Pain: The SENZA-RCT Randomized Controlled Trial. Anesthesiology 123, 851–860. doi: 10.1097/ALN.0000000000000774

45. Keitel, A., and Gross, J. (2016). Individual Human Brain Areas Can Be Identified from Their Characteristic Spectral Activation Fingerprints. PLoS Biol 14, e1002498. doi: 10.1371/journal.pbio.1002498

46. Kim, J. A., Bosma, R. L., Hemington, K. S., Rogachov, A., Osborne, N. R., Cheng, J. C., et al. (2020). Cross-network coupling of neural oscillations in the dynamic pain connectome reflects chronic neuropathic pain in multiple sclerosis. Neuroimage Clin 26, 102230. doi: 10.1016/j.nicl.2020.102230

47. Kirketeig, T., Schultheis, C., Zuidema, X., Hunter, C. W., and Deer, T. (2019). Burst Spinal Cord Stimulation: A Clinical Review. Pain Med 20, S31–S40. doi: 10.1093/pm/pnz003

48. Kong, Q., Li, T., Reddy, S., Hodges, S., and Kong, J. (2024). Brain stimulation targets for chronic pain: Insights from meta-analysis, functional connectivity and literature review. Neurotherapeutics 21, e00297. doi: 10.1016/j.neurot.2023.10.007

49. Lim, M., Jassar, H., Kim, D. J., Nascimento, T. D., and DaSilva, A. F. (2021). Differential alteration of fMRI signal variability in the ascending trigeminal somatosensory and pain modulatory pathways in migraine. J Headache Pain 22, 4. doi: 10.1186/s10194-020-01210-6

50. Lisman, J. E., and Jensen, O. (2013). The Theta-Gamma Neural Code. Neuron 77, 1002–1016. doi: 10.1016/j.neuron.2013.03.007

51. Liu, S., Li, C., Xing, Y., Wang, Y., and Tao, F. (2016). Role of Neuromodulation and Optogenetic Manipulation in Pain Treatment. Curr Neuropharmacol 14, 654–661. doi: 10.2174/1570159X14666160303110503

52. Ma, X., and Khadra, A. (2024). Neural signaling in neuropathic pain: A computational modeling perspective. Current Opinion in Systems Biology 37, 100509. doi: 10.1016/j.coisb.2024.100509

53. Mogedano-Cruz, S., López-Pérez, M., Gijón-Lago, D., Romero-Morales, C., Alonso-Pérez, J. L., Villafañe, J. H., et al. (2025). Peripheral Percutaneous Electrical Nerve Stimulation for Neuropathies: A Systematic Review and Meta-analysis. Pain Manag Nurs 26, 93–101. doi: 10.1016/j.pmn.2024.11.005

54. Näätänen, R., and Picton, T. (1987). The N1 wave of the human electric and magnetic response to sound: a review and an analysis of the component structure. Psychophysiology 24, 375–425. doi: 10.1111/j.1469-8986.1987.tb00311.x

55. Ong, W.-Y., Stohler, C. S., and Herr, D. R. (2019). Role of the Prefrontal Cortex in Pain Processing. Mol Neurobiol 56, 1137–1166. doi: 10.1007/s12035-018-1130-9

56. Oshiro, Y., Quevedo, A. S., McHaffie, J. G., Kraft, R. A., and Coghill, R. C. (2009). Brain mechanisms supporting discrimination of sensory features of pain: a new model. J Neurosci 29, 14924–14931. doi: 10.1523/JNEUROSCI.5538-08.2009

57. Pandya, D. N., and Yeterian, E. H. (1985). “Architecture and Connections of Cortical Association Areas,” in *Association and Auditory Cortices*, eds. A. Peters and E.G. Jones (Boston, MA: Springer US), 3–61. doi: 10.1007/978-1-4757-9619-3_1

58. Paquette, J. P., and Yoo, P. B. (2019). Recruitment of unmyelinated C-fibers mediates the bladder-inhibitory effects of tibial nerve stimulation in a continuous-fill anesthetized rat model. Am J Physiol Renal Physiol 317, F163–F171. doi: 10.1152/ajprenal.00502.2018

59. Parker, T., Huang, Y., Raghu, A. L. B., FitzGerald, J. J., Green, A. L., and Aziz, T. Z. (2020). Dorsal Root Ganglion Stimulation Modulates Cortical Gamma Activity in the Cognitive Dimension of Chronic Pain. Brain Sci 10, 95. doi: 10.3390/brainsci10020095

60. Ploner, M., Sorg, C., and Gross, J. (2017). Brain Rhythms of Pain. Trends Cogn Sci 21, 100–110. doi: 10.1016/j.tics.2016.12.001

61. Puppalla, P., Pattilachan, T. M., Salmeron de Toledo Aguiar, R., Argoff, C. E., and Pilitsis, J. G. (2025). The Future of Pain Management. Neurol Clin 43, 595–615. doi: 10.1016/j.ncl.2025.04.007

62. Ratnadurai-Giridharan, S., Khargonekar, P. P., and Talathi, S. S. (2015). Emergent gamma synchrony in all-to-all interneuronal networks. Front Comput Neurosci 9, 127. doi: 10.3389/fncom.2015.00127

63. Rogers, E. R., Zander, H. J., and Lempka, S. F. (2022). Neural Recruitment During Conventional, Burst, and 10-kHz Spinal Cord Stimulation for Pain. J Pain 23, 434–449. doi: 10.1016/j.jpain.2021.09.005

64. Schulder, M., Mishra, A., Mammis, A., Horn, A., Boutet, A., Blomstedt, P., et al. (2023). Advances in Technical Aspects of Deep Brain Stimulation Surgery. Stereotact Funct Neurosurg 101, 112–134. doi: 10.1159/000529040

65. Sdrulla, A. D., Guan, Y., and Raja, S. N. (2018). Spinal Cord Stimulation: Clinical Efficacy and Potential Mechanisms. Pain Pract 18, 1048–1067. doi: 10.1111/papr.12692

66. Shaheen, N., and Flouty, O. (2024). Unlocking the future of deep brain stimulation: innovations, challenges, and promising horizons. Int J Surg 110, 3146–3148. doi: 10.1097/JS9.0000000000001279

67. Shaheen, N., Shaheen, A., Elgendy, A., Bezchlibnyk, Y. B., Zesiewicz, T., Dalm, B., et al. (2023). Deep brain stimulation for chronic pain: a systematic review and meta-analysis. Front Hum Neurosci 17, 1297894. doi: 10.3389/fnhum.2023.1297894

68. Shirvalkar, P., Sellers, K. K., Schmitgen, A., Prosky, J., Joseph, I., Starr, P. A., et al. (2020). A Deep Brain Stimulation Trial Period for Treating Chronic Pain. J Clin Med 9, 3155. doi: 10.3390/jcm9103155

69. Sirota, A., Montgomery, S., Fujisawa, S., Isomura, Y., Zugaro, M., and Buzsáki, G. (2008). Entrainment of neocortical neurons and gamma oscillations by the hippocampal theta rhythm. Neuron 60, 683–697. doi: 10.1016/j.neuron.2008.09.014

70. Smits, H., Kleef, M. V., Honig, W., Gerver, J., Gobrecht, P., and Joosten, E. A. J. (2009). Spinal cord stimulation induces c-Fos expression in the dorsal horn in rats with neuropathic pain after partial sciatic nerve injury. Neurosci Lett 450, 70–73. doi: 10.1016/j.neulet.2008.11.013

71. Sohal, V. S., Zhang, F., Yizhar, O., and Deisseroth, K. (2009). Parvalbumin neurons and gamma rhythms enhance cortical circuit performance. Nature 459, 698–702. doi: 10.1038/nature07991

72. Song, N., Long, L., Liu, N., Luo, Y., Wei, M., Huang, H., et al. (2025). Harnessing theta waves: tACS as a breakthrough in alleviating post-stroke chronic pain. Front Neurosci 19, 1553862. doi: 10.3389/fnins.2025.1553862

73. Sporns, O., and Betzel, R. F. (2016). Modular Brain Networks. Annu Rev Psychol 67, 613–640. doi: 10.1146/annurev-psych-122414-033634

74. Sun, S., Yin, J., Wei, H., Zeng, Y., Jia, H., and Jin, Y. (2024). Long-Term Efficacy and Safety of High-Frequency Spinal Stimulation for Chronic Pain: A Meta-Analysis of Randomized Controlled Trials. Clin J Pain 40, 415–427. doi: 10.1097/AJP.0000000000001215

75. Taesler, P., and Rose, M. (2016). Prestimulus Theta Oscillations and Connectivity Modulate Pain Perception. J. Neurosci. 36, 5026–5033. doi: 10.1523/JNEUROSCI.3325-15.2016

76. Tan, L. L., and Kuner, R. (2021). Neocortical circuits in pain and pain relief. Nat Rev Neurosci 22, 458–471. doi: 10.1038/s41583-021-00468-2

77. Tan, L. L., Oswald, M. J., Heinl, C., Retana Romero, O. A., Kaushalya, S. K., Monyer, H., et al. (2019). Gamma oscillations in somatosensory cortex recruit prefrontal and descending serotonergic pathways in aversion and nociception. Nat Commun 10, 983. doi: 10.1038/s41467-019-08873-z

78. Thibes, R. B., da Cunha, P. H. M., Lapa, J. D. da S., Dongyang, L., Pinheiro, D. S., Iglesio, R. F., et al. (2025). Intraoperative recordings from the posterior superior insula in awake humans with peripheral neuropathic pain. Neurophysiol Clin 55, 103056. doi: 10.1016/j.neucli.2025.103056

79. Tort, A. B. L., Komorowski, R., Eichenbaum, H., and Kopell, N. (2010). Measuring phase-amplitude coupling between neuronal oscillations of different frequencies. J Neurophysiol 104, 1195–1210. doi: 10.1152/jn.00106.2010

80. Tort, A. B. L., Kramer, M. A., Thorn, C., Gibson, D. J., Kubota, Y., Graybiel, A. M., et al. (2008). Dynamic cross-frequency couplings of local field potential oscillations in rat striatum and hippocampus during performance of a T-maze task. Proc Natl Acad Sci U S A 105, 20517–20522. doi: 10.1073/pnas.0810524105

81. Voytek, B., and Knight, R. T. (2015). Dynamic network communication as a unifying neural basis for cognition, development, aging, and disease. Biol Psychiatry 77, 1089–1097. doi: 10.1016/j.biopsych.2015.04.016

82. Wang, J., and Doan, L. V. (2024). Clinical pain management: Current practice and recent innovations in research. Cell Rep Med 5, 101786. doi: 10.1016/j.xcrm.2024.101786

83. Wang, J., Li, D., Li, X., Liu, F.-Y., Xing, G.-G., Cai, J., et al. (2011). Phase-amplitude coupling between θ and γ oscillations during nociception in rat electroencephalography. Neurosci Lett 499, 84–87. doi: 10.1016/j.neulet.2011.05.037

84. Wang, J., Wang, J., Xing, G.-G., Li, X., and Wan, Y. (2016). Enhanced Gamma Oscillatory Activity in Rats with Chronic Inflammatory Pain. Front Neurosci 10, 489. doi: 10.3389/fnins.2016.00489

85. Zagha, E., Casale, A. E., Sachdev, R. N. S., McGinley, M. J., and McCormick, D. A. (2013). Motor cortex feedback influences sensory processing by modulating network state. Neuron 79, 567. doi: 10.1016/j.neuron.2013.06.008

86. Zhang, T. C., Janik, J. J., and Grill, W. M. (2014). Mechanisms and models of spinal cord stimulation for the treatment of neuropathic pain. Brain Res 1569, 19–31. doi: 10.1016/j.brainres.2014.04.039

