## Supplementary for "High-Frequency Spinal Cord Stimulation Reorganizes Cortical Cross-Frequency Coupling in a Region- and Time-Dependent Manner"

**Supplementary Figures**

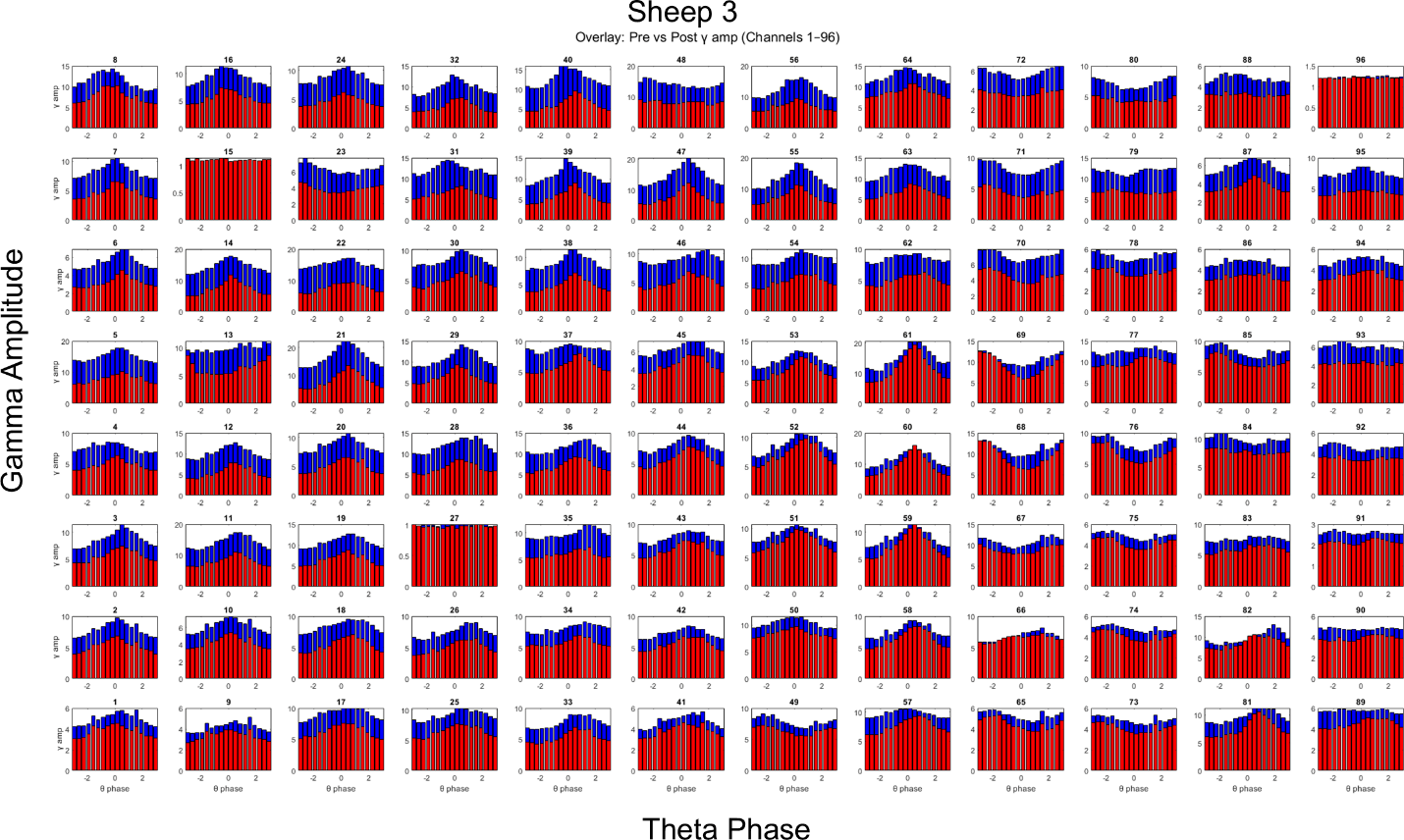

***Supplementary Figure 1****: Theta-phase gamma-amplitude locking for all channels for Sheep 3. Notice an overall decrease in gamma amplitude as reported previously, and similar theta-gamma locking profiles for channels.*

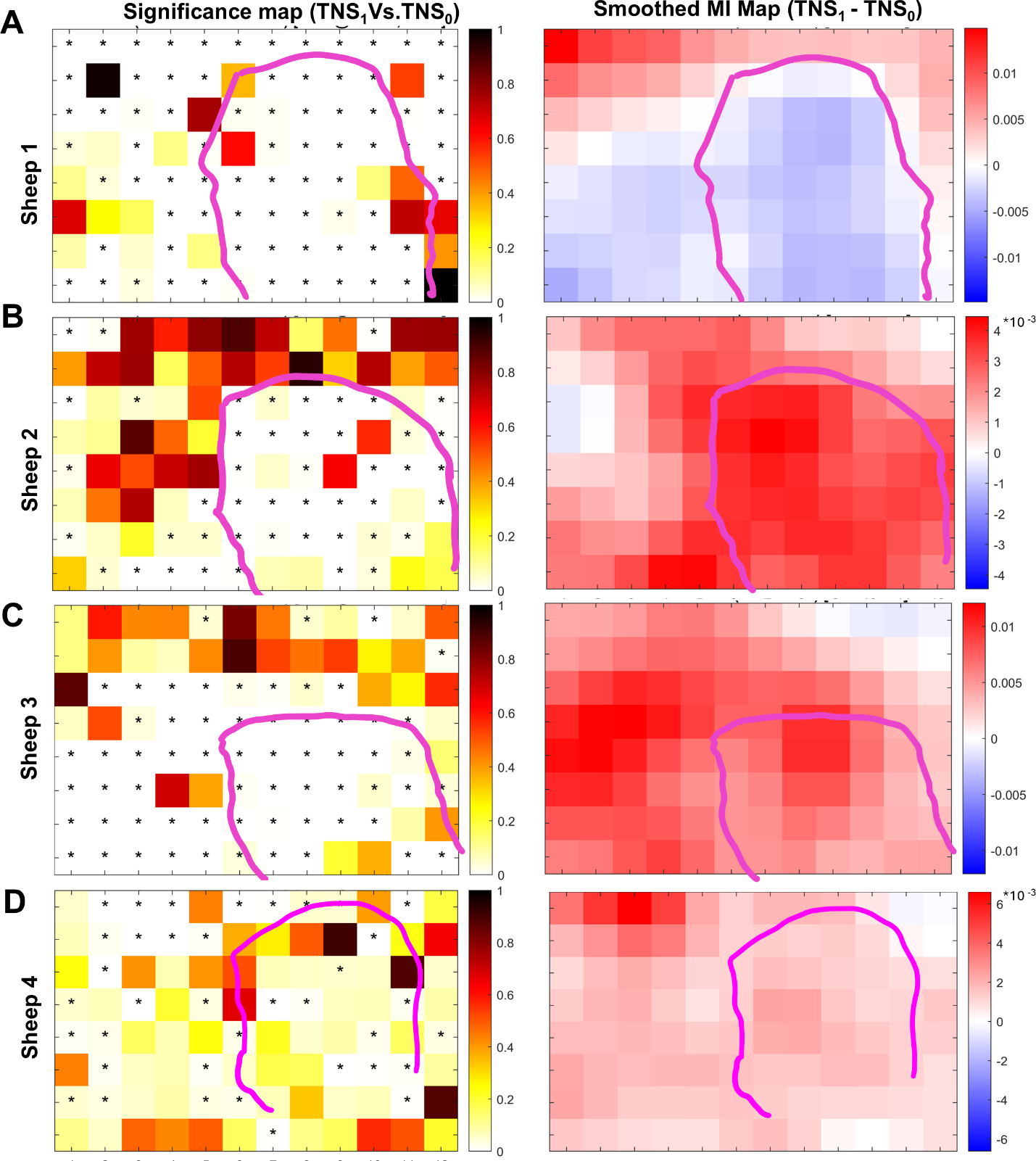

***Supplementary Figure 2****:* ***(A-D****) Left: Channel-by-channel significance (TNS_0_ vs TNS_1_) maps for all sheep (****A-D****: Sheep 1-4, respectively). Right: Subtracted (TNS_1 -_ TNS_0)_ and smoothed MI maps for all sheep. Except Sheep 1, all maps look similar.*

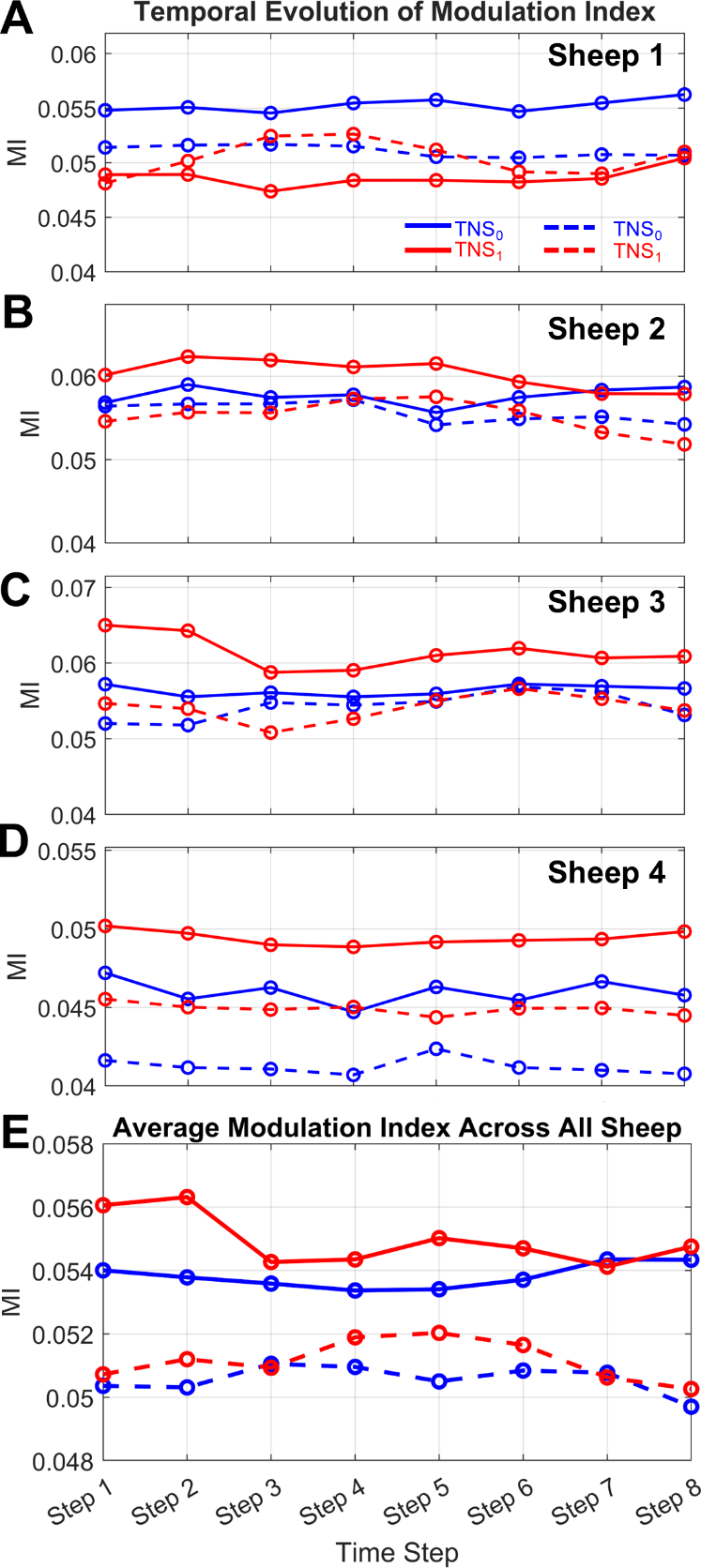

***Supplementary Figure 3:*** *Individual MI trajectories over time for each of the four sheep (****A-D****: Sheep 1-4, respectively), separated by cortical area (association vs. somatosensory) and condition (TNS_0_ vs. TNS_1_). These plots reveal consistent patterns across most animals, except Sheep 1* ***(A)****, which deviates from the group trend. Notably, MI values are higher in the somatosensory cortex than in the association cortex across several sheep, particularly during the early time windows following stimulation.* ***(E)*** *Average MI trajectories show an early peak in the somatosensory cortex (steps 1–2) and a delayed peak in the association cortex (steps 4–6).*
